# Gene-Modulated Network Diffusion for Improved Modeling of Amyloid-*β* Spread in Alzheimer’s Disease

**DOI:** 10.64898/2026.05.04.722725

**Authors:** Frederick H. Xu, Duy Duong-Tran, Heng Huang, Andrew J. Saykin, Paul M. Thompson, Christos Davatzikos, Yize Zhao, Li Shen, the Alzheimer’s Disease Neuroimaging Initiative

**Author notes:** Data used in the preparation of this article were obtained from the Alzheimer’s Disease Neuroimaging Initiative (ADNI) database (adni.loni.usc.edu). As such, the investigators within the ADNI contributed to the design and implementation of ADNI and/or provided data but did not participate in the analysis or writing of this report. A complete listing of ADNI investigators can be found at: http://adni.loni.usc.edu/wp-content/uploads/how_to_apply/ADNI_Acknowledgement_List.pdf.

## Abstract

Understanding the pathogenesis of amyloid-*β* pathology in Alzheimer’s Disease (AD) proves to be a challenge. In this work, we expand upon the application of network diffusion models (NDM) to study pathophysiological spread of amyloid-*β* throughout white matter structural brain networks. We found that the NDM successfully recaptures subpopulation-level spatial patterns (Pearson’s R=0.45-0.48, P_FDR_ *<* 0.01) of amyloid-*β* deposition in the Alzheimer’s Disease Neuroimaging Cohort at a regional level, but with drawbacks in mechanism interpretability. We then moved to an extended NDM framework (eNDM), including a protein synthesis term to better reflect the role of amyloid-*β* metabolism, as well as including regional vulnerability using spatial transcriptomics from the Allen Human Brain Atlas to modulate the region-level rate parameters of the synthesis term. The novel gene eNDMs exhibited significant performance increases in Pearson’s correlation (Steiger’s Z, P_FDR_ *<* 0.10) over baseline NDM performance in mild cognitive impairment and AD groups using APOE, SORL1, and FGL2 for gene modulation. The results were robust and replicable when testing on an external cohort of the Alzheimer’s Disease Sequencing Project. The study thus demonstrates the importance of regional genetic vulnerability, in conjunction with network diffusion mechanisms, in improving the modelling and prediction of amyloid-*β* pathophysiological spread.

## I. Introduction

ALZHEIMER’S disease (AD), an irreversible progressive neurodegenerative disease and the leading cause of dementia, is characterized by a loss of memory recall and cognitive impairment that limits functions of daily living. In the United States, there are currently 6.9 million individuals over the age of 65 afflicted with AD, with the disease now ranking as the 5^th^ leading cause of death in the elderly population as of 2024 [1]. As such, efforts to understand the mechanism of the disease have become ever more important.

A leading hypothesis for the mechanistic progression of AD is the amyloid cascade, whereby the aggregation of amyloid-*β* plaques leads to systemic stress that causes downstream neurodegeneration, neuronal dysfunction, and ultimately cognitive impairment [2], [3]. Early deposition of amyloid has been associated with accelerated development of later pathologies such as tau neurofibrillary tangles or cortical atrophy [4]– [7]. Amyloid load in the brain has thus been considered as a potential early indicator for AD [8], with thresholded amyloid-positivity used in clinical diagnosis [9].

However, while the positive presence and load of amyloid-*β* is highly associated with disease, the cause and course of amyloid spread throughout the brain are not fully understood [10]. Staging studies using post-mortem immunohistochemistry stains [11]–[13] and in vivo studies using positron emission tomography (PET) neuorimaging are able to characterize a spatiotemporal pattern of amyloid spread throughout the brain [14], [15], but are limited in discerning a mechanism for how the pattern is achieved. Genetic sources of abnormality have been associated with amyloid, noting that regional vulnerabilities as identified through spatial transcriptomics offer an elucidating but incomplete explanation for amyloid deposition patterns [16]–[18]. As such, characterizing the pathophysiological progression of amyloid deposition remains a key question in the study of AD, and achieving understanding may enhance efforts for early detection before disease onset with cognitive decline.

One theory to explain the spread of amyloid is the prionlike hypothesis of trans-synaptic spread, which proposes that amyloid propagates through synaptic connections into regions of high vulnerability [19]. Evidence for transsynaptic spread has been found in transgenic mice models, where initial seed injections of amyloid are observed to spread from the seed to interconnected brain regions [20]–[24]. Furthermore, in human studies, trans-synaptic spread has been demonstrated as a viable theory for other proteinopathies in neurodegenerative disease, such as *α*-synuclein in Parkinson’s disease and tau in AD [25]–[27], lending further credence to the theory that amyloid may propagate through a similar mechanism. However, in-vivo exploration of amyloid deposition through neuronal connections in the human brain has remained sparse.

In associated pathologies of cortical atrophy and tau aggregation, studies have found that pathological spread can be explained by diffusion through an underlying brain network structure, where modeling a seed of pathology that then diffuses through the brain network sufficiently recaptures observed spatial patterns of both cortical atrophy and tau aggregation [28], [29]. Further extensions of the network diffusion model and subsequent investigations into regional vulnerability through the lens of spatial transcriptomics have provided key insights into the mechanisms behind tau pathophysiological spread [30]. These advanced neuroimaging techniques bridging between connectomics and pathology offer a unique insight into the potential mechanism of pathogenic spread of proteinopathies through the brain.

To that end, in this work we explore and expand upon a connectivity-driven model of spread to test the hypothesis of trans-synaptic spread of amyloid-*β* through white matter connections. We modeled a seed injection that diffuses through a white matter structural brain network constructed from neuroimaging using a network diffusion model (NDM), and compare the observed distribution of amyloid throughout the network against that of ground truth amyloid positron emission tomography (amyloid-PET) data in three diagnostic groups of healthy control (HC), mild cognitive impairment (MCI), and AD in the Alzheimer’s Disease Neuroimaging Initiative (ADNI) cohort. Our goal is to observe whether a connectivity-driven model of pathogenic spread can plausibly explain observed spatial patterns of amyloid in each subpopulation. A preliminary version of these results has been reported [31]. From there, we investigate extensions of the NDM that better reflect biological hypotheses, including a novel incorporation of regional genetic vulnerabilities using spatial transcriptomics and testing for improvements in model performance. Lastly, we then evaluate the performance of these models on an unseen test cohort to assess model robustness.

## II. Materials and Methods

We extend our work from a previous conference submission [31] that tests NDM performance in the context of amyloid in AD. The present work investigates novel applications of extended network diffusion model frameworks, as well as new methodology to incorporate gene modulation into the model.

### A. Data Acquisition

Data used in the preparation of this article is obtained from the Alzheimer’s Disease Neuroimaging Initiative (ADNI) database (http://adni.loni.usc.edu) [32], [33]. The ADNI was launched in 2003 as a public-private partnership led by Principal Investigator Michael W. Weiner, MD. The primary goal of ADNI has been to test whether serial MRI, PET, other biological markers, and clinical and neuropsychological assessment can be combined to measure the progression of mild cognitive impairment (MCI) and early AD. All participants provided written informed consent, and study protocols were approved by each participating site’s Institutional Review Board (IRB). Up-to-date information about the ADNI is available at www.adni-info.org.

Subjects with available DTI scans, T1-weighted structural MRI (sMRI), and demographic data were obtained from the ADNI-GO/2 database [34]. The subject population consisted of healthy control subjects (n = 34), MCI subjects (n = 55), and AD (n = 28) patients. The sex distribution was 66 male and 55 female subjects. The average age of the total population was 73.0 years with the HC population averaging at 73.2 years, the MCI population averaging at 72.9 years, and the AD population averaging at 73.0 years. We tested for group differences in continuous demographic variables using ANOVA (Table I). No significant difference in ages was detected across population groups (*P* = 0.964). Average number of years of education received was also consistent across population groups (*P* = 0.512), with HC averaging 16.2 years, MCI averaging 15.7 years, and AD averaging 15.4 years. Ethnic distribution in each subject group is predominantly white caucasian (Supplementary Table S1).

**TABLE I:**
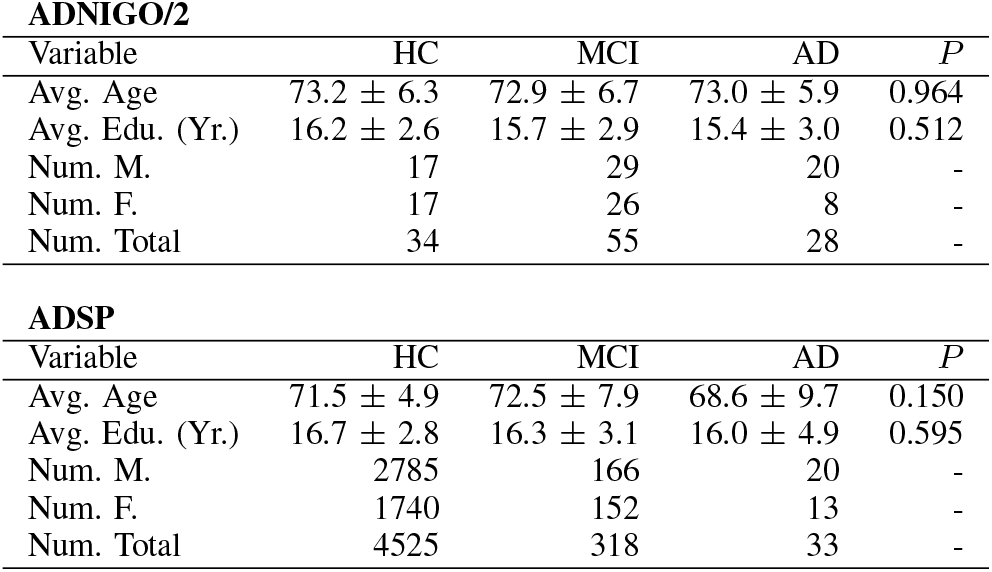
Cohort Population Demographics.

| ADNIGO/2 |  |  |  |  |  |
| --- | --- | --- | --- | --- | --- |
| Variable | HC | MCI | AD | $P$ | |
| Avg. Age | 73.2 ± 6.3 | 72.9 ± 6.7 | 73.0 ± 5.9 | 0.964 | |
| Avg. Edu. (Yr.) | 16.2 ± 2.6 | 15.7 ± 2.9 | 15.4 ± 3.0 | 0.512 | |
| Num. M. | 17 | 29 | 20 | - |  |
| Num. F. | 17 | 26 | 8 | - |  |
| Num. Total | 34 | 55 | 28 | - |  |

| ADSP |  |  |  |  |  |
| --- | --- | --- | --- | --- | --- |
| Variable | HC | MCI | AD | $P$ | |
| Avg. Age | 71.5 ± 4.9 | 72.5 ± 7.9 | 68.6 ± 9.7 | 0.150 | |
| Avg. Edu. (Yr.) | 16.7 ± 2.8 | 16.3 ± 3.1 | 16.0 ± 4.9 | 0.595 | |
| Num. M. | 2785 | 166 | 20 | - |  |
| Num. F. | 1740 | 152 | 13 | - |  |
| Num. Total | 4525 | 318 | 33 | - |  |

### B. Neuroimaging Preprocessing and Network Construction

The DTI data were preprocessed and denoised, motion-corrected, and distortion-corrected using an over-complete local principal components analysis (PCA) [35]. Streamline tractography was performed using a probabilistic algorithm of fiber assignment by continuous tracking (FACT) [36].

sMRI images were then registered to lower resolution b0 volume of the DTI data using the FLIRT toolbox in the FMRIB Software Library (FSL) [37] and 83 cortical and subcortical brain regions of interest (ROIs) were extracted based on the Lausanne 2008 Scale 33 Parcellation [38]. Fiber density networks were then constructed using the procedure from [39], [40]. Each edge in the network’s adjacency matrix **A** is weighted by the fiber density between two ROIs (*i, j*), which is defined as the number of the fibres (NOF) connecting each pair of ROIs divided by their average surface area (SA): **A**(*i, j*) = (NOF_*i,j*_)*/*((SA_*i*_ + SA_*j*_)*/*2) (Figure 1a). Each subjects’ network thus consists of 83 ROI nodes defined by the parcellation, with pairwise measurements of fiber density as the edges between nodes. All edges obtained are retained, without an *a priori* thresholding procedure [41]. For the three subpopulations of HC, MCI, and AD, we average the networks within each subpopulation to obtain group-level averaged networks for study [40], [42].

**Fig. 1.**
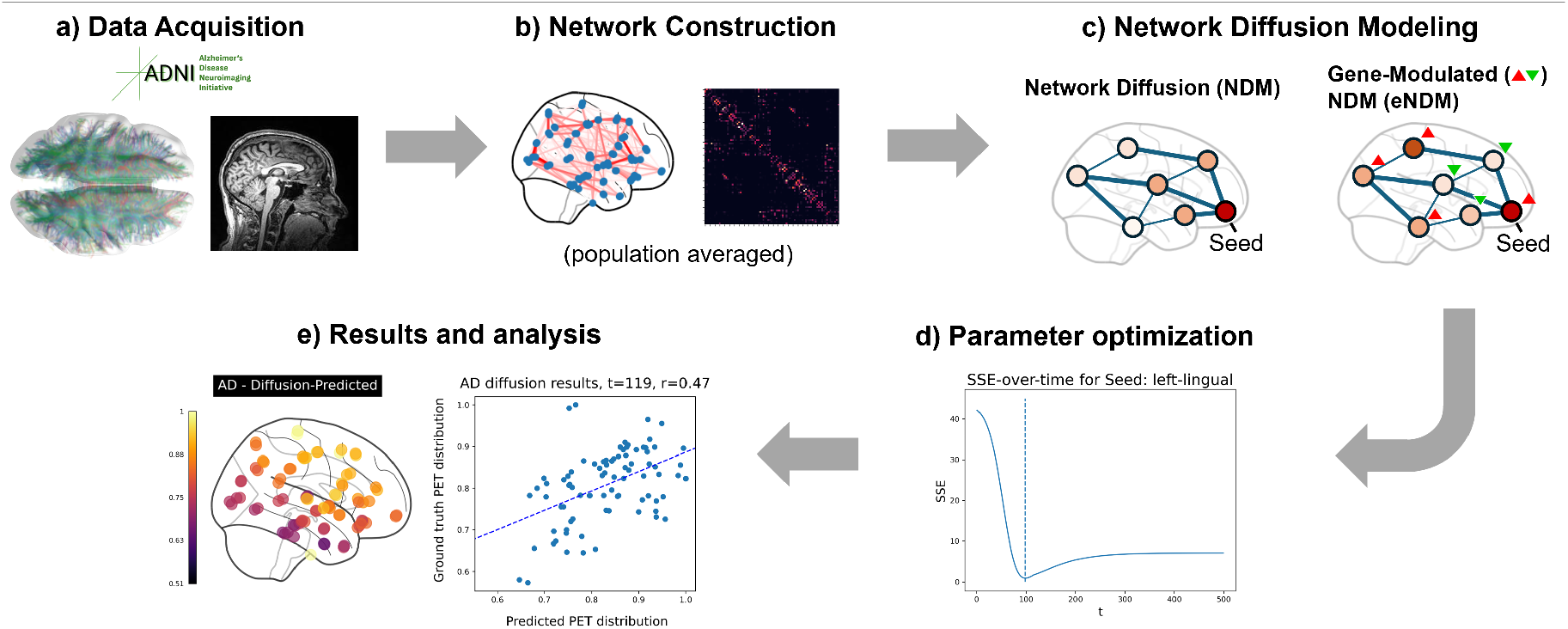
Our main methodology pipeline (a) takes neuroimaging data from the ADNI, (b) processes the data into white matter connectomes, then (c) performs network diffusion to model the spread of pathology from a starting seed, including baseline (NDM) and extended (eNDM) frameworks. (d) We repeat the process, fitting the optimal seed and diffusion time parameters for the NDM by minimizing the SSE between predicted distributions and ground truth distributions observed from FBP PET, and then (e) compare the optimal result against ground truth observations.

### C. Amyloid-β Pathology Data Acquisition and Preprocessing

^18^F-florbetapir (FBP) PET imaging data were then obtained from the ADNI database. For these data, we elected to use the preprocessed 6mm resolution dataset from UC Berkeley, which contains region-level standard uptake value rations (SUVRs) co-registered to an MRI parcellated with the Desikan-Killiany atlas that offers high spatial specificity [43]. To harmonize with our network data, we date-matched the preprocessed FBP PET data to our DTI scans, to the nearest available date within 30 days. Additionally, our parcellation of choice, the Lausanne 2008 Scale 33 parcellation, possesses the same definition of cortical regions as the Desikan-Killiany atlas, thus offering spatial correspondence between the two datasets. We obtained group-level averages of the FBP PET SUVR vectors for each subpopulation of HC, MCI, and AD

For our validation dataset, we chose to use the Alzheimer’s Disease Sequencing Project (ADSP) cohort [44]. The cohort is a consortium of multiple cohorts including the ADNI. Firstly, to avoid data leakage from the training data, we exclude the ADNI data that is present in the ADSP. We then obtain region-level FBP and 18F-florbetaben (FBB) PET SUVRs from the neuroimaging phenotype dataset, registered to and parcellated by the Desikan-Killiany atlas so that the data spatial definition matches that of our training dataset. We note that as part of the ADSP cohort’s data collection procedure, FBP and FBB PET tracers have been harmonized using the centiloid scale [45]. Lastly, we apply ComBat batch harmonization to correct for cohort batch effects, preserving the diagnosis as a covariate [46]. The final size of the ADSP validation dataset is 4876 subjects, with 4525 HC, 318 MCI, and 33 AD subjects. As with the ADNI, there was no significant difference in subpopulation average age or years of education (ANOVA, *P >* 0.05, Table I). The HC population is more male-dominant, while the disease populations are roughly even in sex (Table I).

### D. Region-level genetic expression cohort

Our novel experimental models make use of region-level spatial transcriptomics from the Allen Human Brain Atlas (AHBA). The atlas is derived from 6 healthy subjects. We obtained the microarray genetic expression data that has been registered to the FreeSurfer atlas with 68 cortical regions by previous work in the field [47]. These regions align with 68 cortical regions in our structural brain networks networks.

We then processed the region-level genetic expression into a relative expression value:

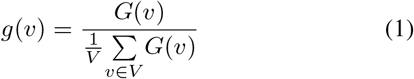

where *g*(*v*) is a vector representing the relative gene expression for a specific gene at node *v* as its nodal gene expression *G*(*v*) divided by its average expression across all nodes (equation 1). If a node possesses above-average gene expression levels, then *g*(*v*) is greater than 1, and vice versa. Using this system, we then imputed relative gene expression values for the subcortical regions that are present in the structural brain networks that lack corresponding genetic expression data in the AHBA dataset. We make the assumption that these regions have average gene expression values, imputing with *g*(*v*_missing_) = 1.

Genes were selected based on prior findings in the literature. APP, PSEN1, and PSEN2 were included due to their close relationship with amyloid-*β* processes and early-onset AD [48]. APOE was also included due to its highly significant statistical relationship with AD and potential role in regulating amyloid-*β* deposition [49]. A group of genes (CLU, PICALM, CR1, BIN1, ABCA7, SORL1) identified through a review of AD genome-wide association (GWAS) studies [50] was included, as well as a group (TOMM40, FGL2, IAPP, SLCO1A2) from a review of amyloid PET-GWAS studies [51]. Gene selection was subject to availability in the AHBA cohort.

### E. Baseline Network Diffusion Model

To model the spread of amyloid through a network backbone, we turn to the theory of heat diffusion through space. This theory has been applied in previous work in the field to structural brain networks to observe the spread of cortical atrophy and tau proteins [28], [29]. We assume propagation of amyloid between connected brain regions described by the network heat equation:

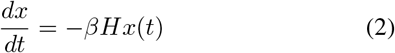

where the vector *x*(*t*) is the distribution of pathology in each region at time *t, β* is the diffusivity constant, and **H** is the Laplacian of the adjacency matrix **A** of the structural brain network connectome.

These definitions arrive at the traditional network diffusion model (NDM), which we treat as the baseline to compare novel experimental models against.

### F. Extended Network Diffusion Model

We introduce extended variants of the NDM using a generalized extended network diffusion model (eNDM) framework:

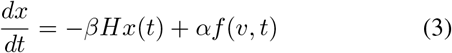

where an additional synthesis term *αf* (*v, t*) denotes a node-level process occurring in the space of nodes *v ∈ V* defined by the Lausanne Parcellation, time *t*, and controlled by *α* which governs the importance [30]. The function *f* (*v, t*) can thus take on various definitions for novel experimental eNDMs.

1. *Seed-based Kinetics:* Previous work has proposed the following definition for an extended network diffusion model:

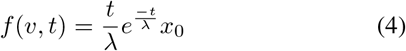

where *f* (*v, t*) takes the form of a gamma function to model first-order synthesis kinetics for proteinopathy agents occurring with rate *λ* at the seed defined through the initialization *x*_0_ [30]. While this model presents an early iteration of eNDMs, it make a significant assumption that the synthesis only occurs at a pathological epicenter. This assumption is at odds with currently literature hypotheses on amyloid-*β* deposition where the phenomenon is suggested to be more widespread with multi-regional susceptibilities [52], [53].
2. *Global Kinetics:* We thus propose novel definitions of eNDMs. The first is extending the kinetics term to apply to all nodes in the network instead of just the seed. Modifying equation 4, we extend to all nodes:

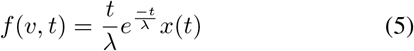

where the gamma function is multiplied with *x*(*t*). Recall that *x*(*t*) is a vector of length *V* that denotes the regional distribution of pathology at time *t*. The end result models an amyloid cascade hypothesis: if the pathology distribution is high in a region due to passive diffusion, then the kinetics term at that region is amplified, thus boosting the importance of pathology synthesis.
3. *Gene-modulated Kinetics:* To further extend the model in equation 5, we introduce a regional susceptibility measure determined by gene expression at each node:

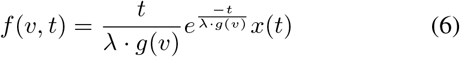

where the rate parameter *λ* is modulated by a relative gene expression value *g*(*v*) at each node for a specific gene *G* (equation 1). If a node *u* possesses above-average gene expression levels, then *g*(*u*) *>* 1 and *λ* is amplified, and vice versa. We thus arrive at the gene-modulated extended network diffusion model (gene eNDM).

### G. Numerical Solution for NDMs

While the baseline network diffusion model (equation 2) has a closed-form solution [28], we chose to solve the differential equations numerically using the finite difference method so the same procedure could be applied to the eNDM framework (equation 3) where a closed-form solution is unavailable.

In-line with previous work in the field, the initial distribution *x*_0_ at *t* = 0 is seeded with a value of 1 at the seed region and 0 elsewhere [29]. Diffusion time *t* is then iterated using a time step and *x*(*t*) is calculated using the finite difference method. The distribution *x*(*t*) is then normalized at each step such that the values fall in the range [0, 1].

### H. Model Optimization and Evaluation

To find the optimal parameters for the network diffusion model, minimize the sum of squared errors (SSE) between the predicted distribution *x*(*t*) and the ground truth distribution *y*(*t*) of FBP PET SUVR values. In line with previous work in the field, we hold *β* constant. We chose a value of *β* = 0.0025 for all models. The remaining parameters of the respective models are then optimized by a gridsearch.

The baseline NDM offers two potential parameters of optimization. Firstly, we may tune the diffusion time *t* by holding the diffusivity constant *β* constant. As such, we iterate through *t* = [0, 300] with a time step *t*_step_ = 1 to find the optimal diffusion time. The second parameter that may be tuned is the starting seed. We thus iterate through each ROI to serve as a starting seed to find the optimal choice of seed. We select the optimal combination of *t* and starting seed based on the lowest SSE (Figure 1d). The eNDM framework offers 4 potential parameters of optimization: diffusion time *t*, the optimal starting seed, the weighting *α* of the kinetic function *f* (*v, t*), and the rate parameter *λ*. We optimize within a range of *t* = [0, 300] with *t*_step_ = 5, *α* = (0, 2) in steps of 0.2, *λ* = (20, 150) in steps of 10, and with each of the 83 ROIs as a potential starting seed in the gridsearch. We note that the time step is increased here to reduce the computational burden of fitting an eNDM for each individual gene. The optimal combination of these parameters is chosen through minimizing SSE between predicted *x*(*t*) and ground truth *y*(*t*) pathology distributions for each eNDM.

We evaluated the model using the Pearson correlation (*R*) between *x*(*t*) and *y*(*t*), as well as the *P* -value to assess the significance of the correlation. We apply a false discovery rate (FDR) correction to correct for multiple comparisons. For eNDMs, we computed the performance improvement over baseline by evaluating the difference in Pearson’s correlation between the experimental eNDMs and the baseline NDM (Δ*R* = *R*_eNDM_ − *R*_NDM_). To assess whether the improvement in model performance is significant, we conducted a one-sided Steiger’s Z-test for differences in dependent correlations, with a null hypothesis that the models do not offer improvements in performance. This test was chosen because both correlations are evaluated with respect to a common baseline ground truth, and thus share a dependency. *P* -values from this test were then corrected for multiple comparisons using FDR correction.

### I. Model Validation on External Cohort

After fitting each model using the ADNI data, we sought to test model performance on an unseen ADSP cohort dataset. We froze eNDM parameters and used the models and structural brain network data from ADNI to predict on the regional amyloid PET pathology data from ADSP. We report the performance metrics of Pearson’s R, significance of Pearson’s R, SSE, which are computed from comparing the model prediction against the ADSP regional PET as ground truth. We also report difference-from-baseline metrics comparing eNDM prediction performance on ADSP against the baseline NDM performance on ADSP.

## III. Results

### A. Population-level Network Diffusion

We first fit an NDM at the diagnostic subpopulation-level for the ADNI GO/2 cohort, using averaged structural brain networks and averaged regional PET SUVRs. Predicted PET distributions within each subpopulation achieved statistically significant fits (*P <* 0.01), with Pearson’s correlation between prediction and ground truth ranging from 0.44-0.48 for all 3 subpopulations (Table II, Figure 3). The optimal diffusion time (*t*) was found to increase monotonically with disease severity (*t*_HC_ = 98 *< t*_MCI_ = 104 *< t*_AD_ = 119). For the optimal seed, it was found that the brainstem was the optimal seeding region for all 3 of the subpopulations. When qualitatively observing the spatial pattern of the PET distribution, we conclude that the spatial pattern was adequately recaptured (Figure 3). The NDM results are presented from previous work [31] and serve as a baseline model for subsequent experimental eNDMs to compare against.

**TABLE II:** ADNI Network Diffusion Model Result Metrics.

| Model | Gene | Group | $R$ | $P$-val | $P_{FDR}$ | SSE | $R^2$ | $\Delta R$ | Steiger-Z $P$ | Steiger-Z $P_{FDR}$ |
| --- | --- | --- | --- | --- | --- | --- | --- | --- | --- | --- |
| NDM | | HC | 0.451 | $1.9 \times 10^{-5}$ | $3.7 \times 10^{-5}$ | 0.976 | 0.204 | 0 | N/A | N/A |
| NDM | | MCI | 0.449 | $2.1 \times 10^{-5}$ | $4.0 \times 10^{-5}$ | 0.821 | 0.201 | 0 | N/A | N/A |
| NDM | | AD | 0.473 | $6.2 \times 10^{-6}$ | $1.6 \times 10^{-5}$ | 0.713 | 0.224 | 0 | N/A | N/A |
| eNDM-seed | | HC | 0.496 | $1.9 \times 10^{-6}$ | $1.1 \times 10^{-5}$ | 0.453 | 0.246 | 0.045 | 0.10 | 0.24 |
| eNDM-seed | | MCI | 0.448 | $2.2 \times 10^{-5}$ | $4.1 \times 10^{-5}$ | 0.449 | 0.200 | -0.001 | 0.49 | 0.49 |
| eNDM-seed | | AD | 0.474 | $6.1 \times 10^{-6}$ | $1.6 \times 10^{-5}$ | 0.571 | 0.224 | 0.0003 | 0.49 | 0.49 |
| eNDM-global | | HC | 0.451 | $1.9 \times 10^{-5}$ | $3.7 \times 10^{-5}$ | 0.976 | 0.204 | 0 | N/A | N/A |
| eNDM-global | | MCI | 0.449 | $2.1 \times 10^{-5}$ | $4.0 \times 10^{-5}$ | 0.821 | 0.201 | 0 | N/A | N/A |
| eNDM-global | | AD | 0.473 | $6.2 \times 10^{-6}$ | $1.6 \times 10^{-5}$ | 0.713 | 0.224 | 0 | N/A | N/A |
| eNDM_gene | APP | HC | 0.386 | $3.1 \times 10^{-4}$ | $3.7 \times 10^{-4}$ | 0.895 | 0.149 | -0.065 | 0.11 | 0.24 |
| eNDM_gene | APP | MCI | 0.436 | $3.8 \times 10^{-5}$ | $5.5 \times 10^{-5}$ | 0.704 | 0.190 | -0.013 | 0.40 | 0.48 |
| eNDM_gene | APP | AD | 0.469 | $7.9 \times 10^{-6}$ | $1.8 \times 10^{-5}$ | 0.624 | 0.220 | -0.005 | 0.46 | 0.49 |
| eNDM_gene | PSEN1 | HC | 0.415 | $9.7 \times 10^{-5}$ | $1.2 \times 10^{-4}$ | 0.743 | 0.172 | -0.036 | 0.26 | 0.40 |
| eNDM_gene | PSEN1 | MCI | 0.356 | $9.7 \times 10^{-4}$ | 0.0011 | 0.632 | 0.126 | -0.093 | 0.16 | 0.33 |
| eNDM_gene | PSEN1 | AD | 0.510 | $8.3 \times 10^{-7}$ | $1.1 \times 10^{-5}$ | 0.549 | 0.260 | 0.037 | 0.27 | 0.40 |
| eNDM_gene | PSEN2 | HC | 0.444 | $2.6 \times 10^{-5}$ | $4.4 \times 10^{-5}$ | 1.244 | 0.197 | -0.007 | 0.089 | 0.23 |
| eNDM_gene | PSEN2 | MCI | 0.428 | $5.3 \times 10^{-5}$ | $7.1 \times 10^{-5}$ | 0.853 | 0.184 | -0.020 | 0.035 | 0.11 |
| eNDM_gene | PSEN2 | AD | 0.476 | $5.5 \times 10^{-6}$ | $1.6 \times 10^{-5}$ | 0.674 | 0.226 | 0.002 | 0.47 | 0.49 |
| eNDM_gene | APOE | HC | -0.204 | 0.064 | 0.065 | 0.944 | 0.042 | -0.656 | $5.7 \times 10^{-6}$ | $8.5 \times 10^{-5}$ |
| eNDM_gene | APOE | MCI | -0.262 | 0.017 | 0.017 | 0.870 | 0.068 | -0.710 | $1.4 \times 10^{-7}$ | $6.3 \times 10^{-6}$ |
| eNDM_gene | APOE | AD | 0.500 | $1.5 \times 10^{-6}$ | $1.1 \times 10^{-5}$ | 0.699 | 0.250 | 0.027 | $1.5 \times 10^{-4}$ | 0.0016 |
| eNDM_gene | SORL1 | HC | 0.458 | $1.3 \times 10^{-5}$ | $2.9 \times 10^{-5}$ | 1.243 | 0.210 | 0.007 | 0.44 | 0.49 |
| eNDM_gene | SORL1 | MCI | 0.547 | $9.0 \times 10^{-8}$ | $4.3 \times 10^{-6}$ | 0.699 | 0.299 | 0.098 | 0.020 | 0.08 |
| eNDM_gene | SORL1 | AD | 0.508 | $9.5 \times 10^{-7}$ | $1.1 \times 10^{-5}$ | 0.682 | 0.258 | 0.035 | 0.015 | 0.068 |
| eNDM_gene | CLU | HC | 0.447 | $2.3 \times 10^{-5}$ | $4.1 \times 10^{-5}$ | 1.200 | 0.199 | -0.005 | 0.28 | 0.40 |
| eNDM_gene | CLU | MCI | 0.453 | $1.7 \times 10^{-5}$ | $3.5 \times 10^{-5}$ | 0.882 | 0.206 | 0.005 | 0.23 | 0.40 |
| eNDM_gene | CLU | AD | 0.484 | $3.6 \times 10^{-6}$ | $1.4 \times 10^{-5}$ | 0.711 | 0.234 | 0.010 | 0.033 | 0.11 |
| eNDM_gene | PICALM | HC | 0.444 | $2.7 \times 10^{-5}$ | $4.4 \times 10^{-5}$ | 1.120 | 0.197 | -0.008 | 0.40 | 0.48 |
| eNDM_gene | PICALM | MCI | 0.496 | $1.8 \times 10^{-6}$ | $1.1 \times 10^{-5}$ | 0.811 | 0.246 | 0.048 | 0.092 | 0.23 |
| eNDM_gene | PICALM | AD | 0.516 | $5.9 \times 10^{-7}$ | $1.1 \times 10^{-5}$ | 0.641 | 0.266 | 0.043 | 0.038 | 0.11 |
| eNDM_gene | CR1 | HC | 0.320 | 0.0031 | 0.0033 | 1.127 | 0.103 | -0.131 | 0.006 | 0.035 |
| eNDM_gene | CR1 | MCI | 0.468 | $8.0 \times 10^{-6}$ | $1.8 \times 10^{-5}$ | 0.722 | 0.219 | 0.020 | 0.26 | 0.40 |
| eNDM_gene | CR1 | AD | 0.479 | $4.7 \times 10^{-6}$ | $1.5 \times 10^{-5}$ | 0.695 | 0.229 | 0.005 | 0.43 | 0.49 |
| eNDM_gene | BIN1 | HC | 0.438 | $3.4 \times 10^{-5}$ | $5.1 \times 10^{-5}$ | 1.020 | 0.192 | -0.013 | 0.39 | 0.48 |
| eNDM_gene | BIN1 | MCI | 0.428 | $5.3 \times 10^{-5}$ | $7.1 \times 10^{-5}$ | 0.711 | 0.184 | -0.020 | 0.31 | 0.43 |
| eNDM_gene | BIN1 | AD | 0.490 | $2.5 \times 10^{-6}$ | $1.3 \times 10^{-5}$ | 0.623 | 0.240 | 0.017 | 0.30 | 0.42 |
| eNDM_gene | ABCA7 | HC | 0.394 | $2.3 \times 10^{-4}$ | $2.7 \times 10^{-4}$ | 0.978 | 0.155 | -0.057 | 0.067 | 0.19 |
| eNDM_gene | ABCA7 | MCI | 0.421 | $7.5 \times 10^{-5}$ | $9.7 \times 10^{-5}$ | 0.793 | 0.177 | -0.028 | 0.19 | 0.35 |
| eNDM_gene | ABCA7 | AD | 0.485 | $3.4 \times 10^{-6}$ | $1.4 \times 10^{-5}$ | 0.667 | 0.235 | 0.011 | 0.27 | 0.40 |
| eNDM_gene | TOMM40 | HC | 0.350 | 0.0011 | 0.0013 | 1.145 | 0.122 | -0.101 | 0.035 | 0.11 |
| eNDM_gene | TOMM40 | MCI | 0.438 | $3.4 \times 10^{-5}$ | $5.1 \times 10^{-5}$ | 0.859 | 0.192 | -0.010 | 0.32 | 0.43 |
| eNDM_gene | TOMM40 | AD | 0.479 | $4.7 \times 10^{-6}$ | $1.5 \times 10^{-5}$ | 0.722 | 0.229 | 0.005 | 0.22 | 0.39 |
| eNDM_gene | FGL2 | HC | -0.267 | 0.015 | 0.015 | 1.169 | 0.071 | -0.718 | $1.57 \times 10^{-6}$ | $3.5 \times 10^{-5}$ |
| eNDM_gene | FGL2 | MCI | 0.478 | $5.0 \times 10^{-6}$ | $1.5 \times 10^{-5}$ | 0.766 | 0.228 | 0.029 | 0.0058 | 0.035 |
| eNDM_gene | FGL2 | AD | 0.499 | $1.6 \times 10^{-6}$ | $1.1 \times 10^{-5}$ | 0.691 | 0.249 | 0.026 | 0.0033 | 0.030 |
| eNDM_gene | IAPP | HC | 0.409 | $1.2 \times 10^{-4}$ | $1.5 \times 10^{-4}$ | 1.208 | 0.168 | -0.042 | 0.015 | 0.068 |
| eNDM_gene | IAPP | MCI | 0.443 | $2.7 \times 10^{-5}$ | $4.4 \times 10^{-5}$ | 0.891 | 0.196 | -0.0056 | 0.18 | 0.35 |
| eNDM_gene | IAPP | AD | 0.489 | $2.8 \times 10^{-6}$ | $1.3 \times 10^{-5}$ | 0.709 | 0.239 | 0.015 | 0.0055 | 0.035 |
| eNDM_gene | SLCO1A2 | HC | 0.384 | $3.4 \times 10^{-4}$ | $3.9 \times 10^{-4}$ | 0.800 | 0.148 | -0.067 | 0.14 | 0.30 |
| eNDM_gene | SLCO1A2 | MCI | 0.429 | $5.2 \times 10^{-5}$ | $7.1 \times 10^{-5}$ | 0.686 | 0.184 | -0.020 | 0.37 | 0.48 |
| eNDM_gene | SLCO1A2 | AD | 0.477 | $5.0 \times 10^{-6}$ | $1.5 \times 10^{-5}$ | 0.607 | 0.228 | 0.004 | 0.48 | 0.49 |

### B. Seed Kinetics

We then tested definitions of eNDMs from previous work in the field, applying a kinetic synthesis term at the seed region and fitting the models at a subpopulation level. Performance was found to be similar to the baseline NDM with significant Pearson’s correlations ranging from 0.45-0.49 (*P <* 0.01, Table II) and a lower SSE than the baseline NDM. However, we observed qualitatively that the seed kinetics eNDM exhibited more clustering of values in the lower-proportion regime of the distribution, with a strong outlier at the seed region in all three diagnostic groups (Figure 2). As such, the performance is similar to the baseline model (Steiger Z-test *P >* 0.05, Table II) but significant room for improvement.

**Fig. 2.**
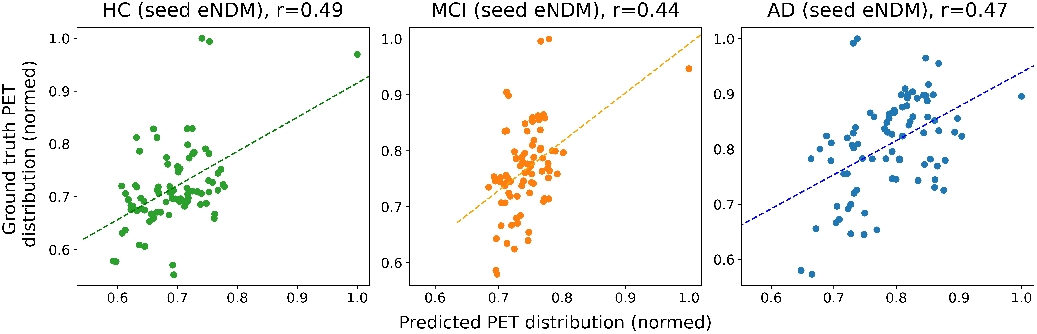
Performance of the eNDM with a kinetic term application at the seed.

**Fig. 3.**
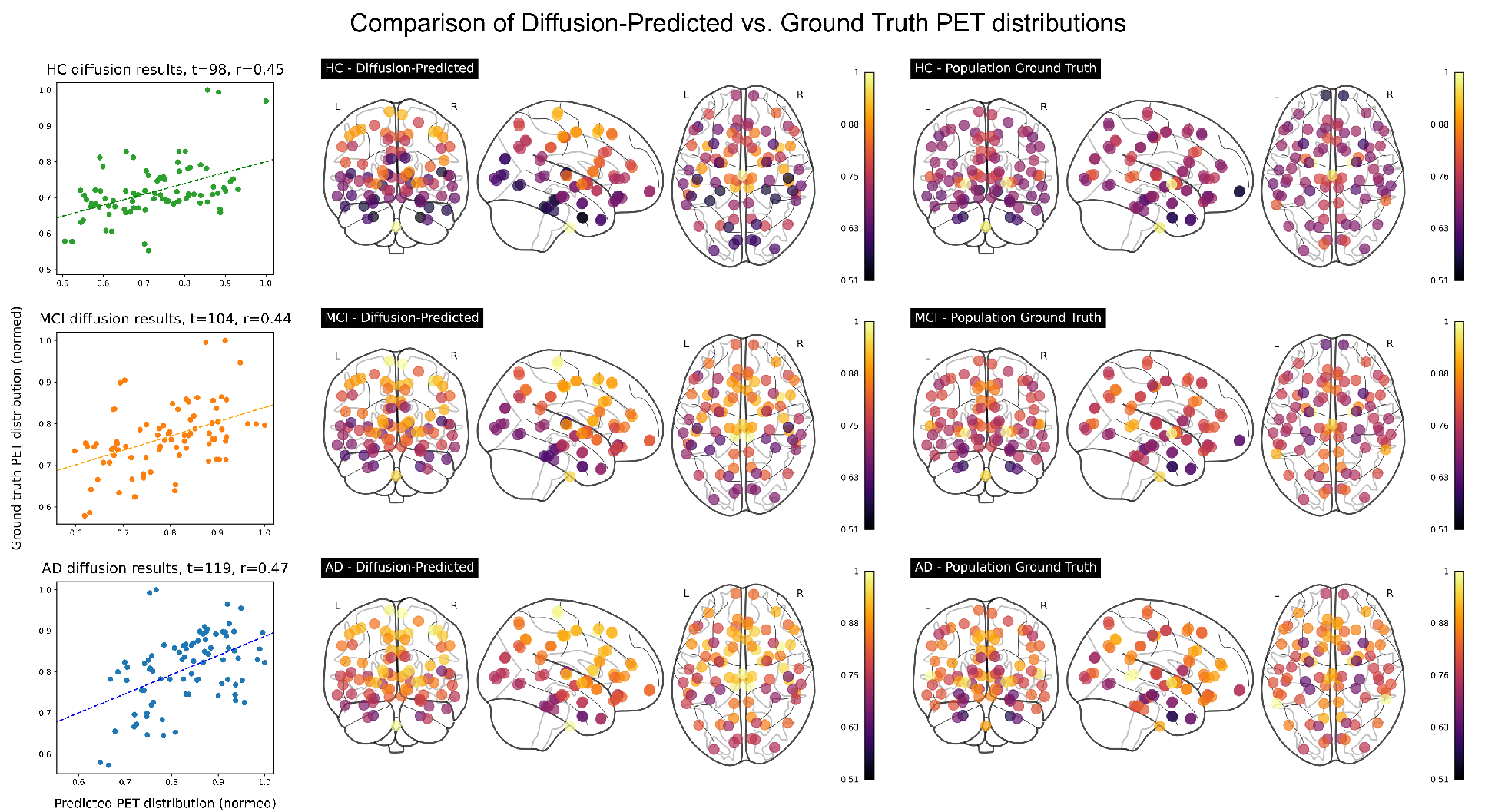
A comparison of amyloid-PET distribution predicted using the baseline NDM, compared against the subpopulation-average ground truth distribution computed from averaged FBP PET SUVR values within each diagnostic group.

### C. Global Kinetics Do Not Improve Model Performance

We then applied the kinetics term to all regions *x*(*t*) instead of only at the seed *x*_0_, thus modeling a wide-spread simultaneous effect of amyloid synthesis. We found that applying the term globally did not result in performance increases. Pearson’s *R* remained the same in all three groups compared to that of the baseline NDM (Table II). The diffusion time *t* was observed to increase (Supplementary Table S2), but no other impact on model performance was observed.

### D. Gene-modulation Impact on eNDM Performance

We then modulated the kinetics term using a region-level vulnerability, identified through regional gene expression values in the AHBA cohort, and fit a GM-eNDM model for each gene using ADNI network and pathology data. Comparing performance gain in Pearson’s *R* using a 1-sided Steiger’s z-test, we found that three genes exhibited gains in disease groups (*P <* 0.05, FDR-corrected). APOE and IAPP improved AD group model performance significantly, while FGL2 improved both MCI group and AD group model performances (Table II). When loosening the significance threshold slightly to *P <* 0.10, we found SORL1 offered significant gains in MCI and AD. Gain magnitude of SORL1 was also the highest among the genes with significant improvements, with Δ*R* = 0.098 in the MCI group and Δ*R* = 0.034 in the AD group (Table II, Figure 4B). Qualitatively, we observe that the 4 genes offer better-resolved AD group distributions with more even distributions that better match ground truth (Figure 4A). Additionally, we qualitatively observe that SORL1 offers significantly improved prediction in MCI compared to the other models. As such, we have identified a group of genes that enhance the network diffusion model, thus explaining a larger proportion of the variance in ground truth amyloid data. We also observed that gene model performance in HC generally did not reject the null hypothesis, often resulting in worsened performance when introducing gene modulation (Table II, Figure 4C). In particular, we note that APOE and FGL2 modulation resulted in anticorrelations with ground truth in the HC group (Table II, Figure 4A).

**Fig. 4.**
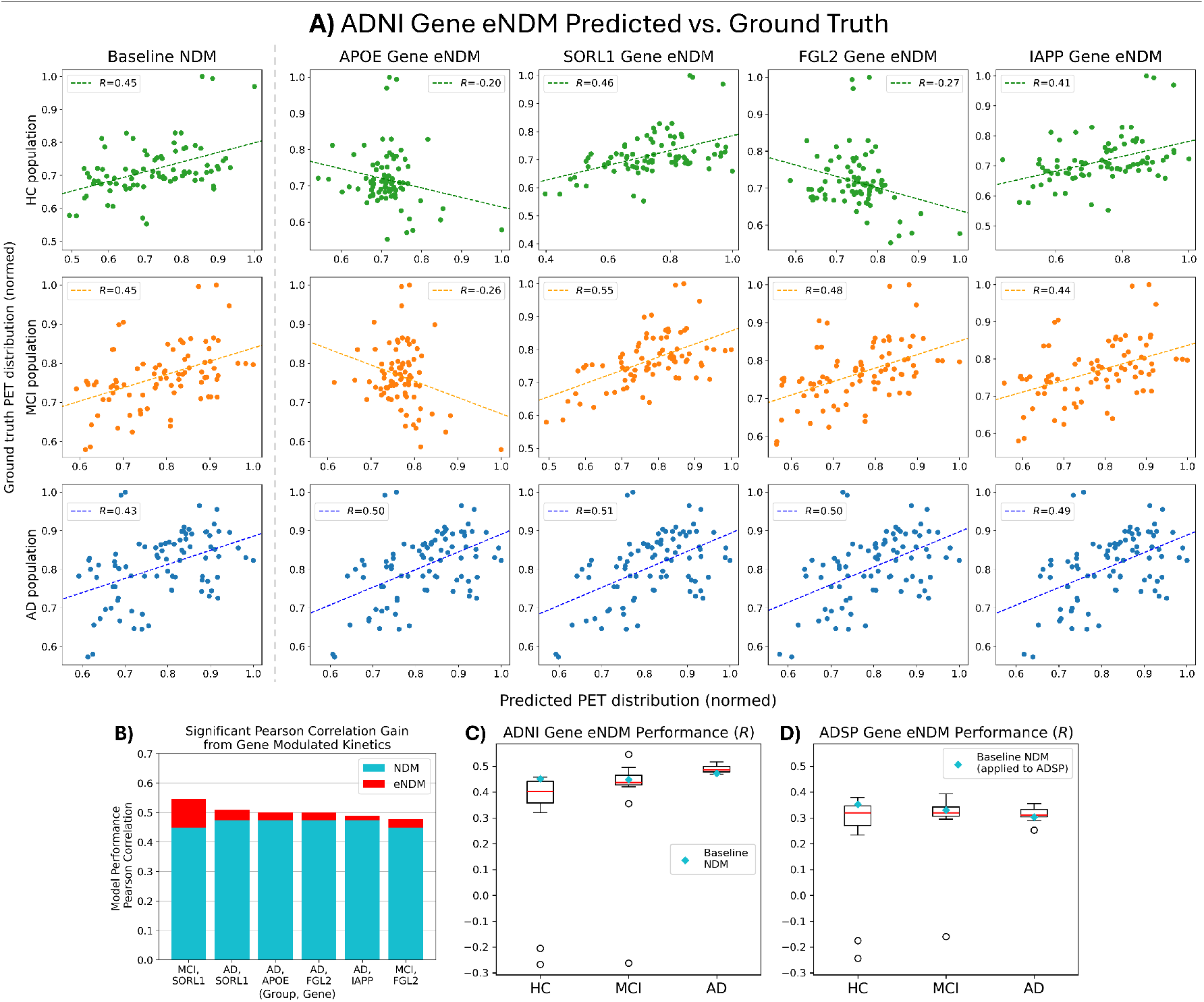
(A) A comparison of amyloid-PET distribution predicted using the gene NDMs, plotting predicted distribution against the ground truth with a line-of-best-fit, depicted for genes with significant improvements over baseline (Steiger’s Z-test, ***P***_FDR_ ***<* 0.10**) in one or more diagnostic groups. The baseline NDM performance plots (left) are provided for qualitative comparisons. (B) Bar plot depicts significant **Δ*R*** gain (Steiger’s Z-test ***P***_FDR_ ***<* 0.10**), ordered by magnitude, plotted against the baseline NDM performance. (C) Boxplot showing the distribution of all ADNI gene eNDM performance ***R*** values in each subpopulation, with a red diamond for baseline NDM performance in ADNI. (D) We depict a similar plot as in (C) for the ADSP cohort gene eNDM results.

### E. ADSP Validation Confirms Results

When testing model generalizability on predicting an unseen cohort’s regional amyloid PET pathology, we observed the NDM was successfully able to predict the spatial distribution of pathology with moderate but significant Pearson’s correlations (Pearson’s *R*=0.30-0.38, *P <* 0.01). Testing the eNDMs on the ADSP revealed similar results as in the ADNI. Most notably, we observed significant performance improvements (Steiger’s Z, *P*_FDR_ *<* 0.05, Table III) in the ADSP AD group for the APOE, SORL1, and FGL2 gene eNDMs, aligning with findings in the ADNI cohort. Model performances are overall weaker in each of the respective subpopulations compared to model performances in the ADNI (Figure 4), but still remain largely significant (Pearson’s *R, P <* 0.01, Supplementary Table S3). These models thus demonstrate good generalizability, as well as elucidating significant and stable contributions from genes in aiding eNDM performance.

**TABLE III:**
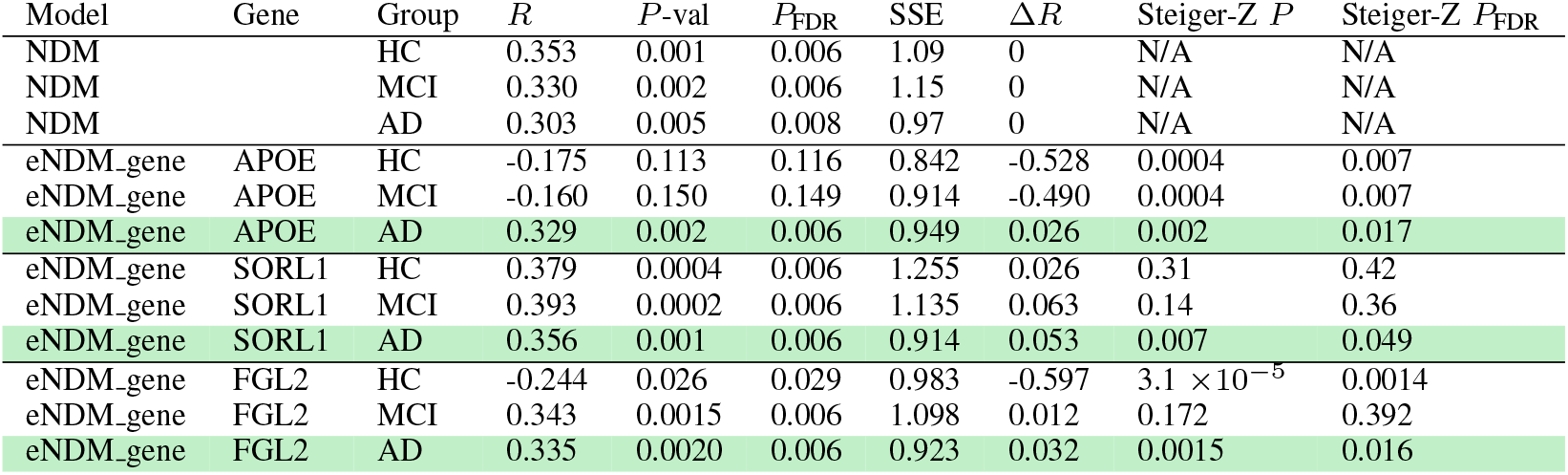
ADSP Validation Network Diffusion Model Result Metrics - Genes With Significant Increases Over Baseline.

| Model | Gene | Group | $R$ | $P$-val | $P_{FDR}$ | SSE | $\Delta R$ | Steiger-Z $P$ | Steiger-Z $P_{FDR}$ |
| --- | --- | --- | --- | --- | --- | --- | --- | --- | --- |
| NDM |  | HC | 0.353 | 0.001 | 0.006 | 1.09 | 0 | N/A | N/A |
| NDM |  | MCI | 0.330 | 0.002 | 0.006 | 1.15 | 0 | N/A | N/A |
| NDM |  | AD | 0.303 | 0.005 | 0.008 | 0.97 | 0 | N/A | N/A |
| eNDM_gene | APOE | HC | -0.175 | 0.113 | 0.116 | 0.842 | -0.528 | 0.0004 | 0.007 |
| eNDM_gene | APOE | MCI | -0.160 | 0.150 | 0.149 | 0.914 | -0.490 | 0.0004 | 0.007 |
| eNDM_gene | APOE | AD | 0.329 | 0.002 | 0.006 | 0.949 | 0.026 | 0.002 | 0.017 |
| eNDM_gene | SORL1 | HC | 0.379 | 0.0004 | 0.006 | 1.255 | 0.026 | 0.31 | 0.42 |
| eNDM_gene | SORL1 | MCI | 0.393 | 0.0002 | 0.006 | 1.135 | 0.063 | 0.14 | 0.36 |
| eNDM_gene | SORL1 | AD | 0.356 | 0.001 | 0.006 | 0.914 | 0.053 | 0.007 | 0.049 |
| eNDM_gene | FGL2 | HC | -0.244 | 0.026 | 0.029 | 0.983 | -0.597 | $3.1 \times 10^{-5}$ | 0.0014 |
| eNDM_gene | FGL2 | MCI | 0.343 | 0.0015 | 0.006 | 1.098 | 0.012 | 0.172 | 0.392 |
| eNDM_gene | FGL2 | AD | 0.335 | 0.0020 | 0.006 | 0.923 | 0.032 | 0.0015 | 0.016 |

### V. Discussion

In this work, we tested established and novel network diffusion models to see whether the distribution of amyloid in the brain in HC, MCI, and AD subpopulations of the ADNI could be modeled through a network diffusion process on a white matter structural brain network backbone. Our previous findings show that the regional amyloid distributions can be adequately recaptured at the population level, with statistically significant fits and correlation values of 0.44-0.48 between predicted distributions and ground truth FBP PET SUVR distributions found using NDMs [31].

Previous work has demonstrated that the passive diffusion mechanism of the NDM occurs through a white matter fiber density network backbone derived from tractography of neuronal tracts identified through DTI, and that the network topology plays a significant role in the process. The findings thus provided evidence for the trans-synaptic spread hypothesis of amyloid [19], where disease agents are hypothesized to propagate through neurons and synaptic connections in a prion-like fashion. The significant involvement of the white matter brain network in the diffusion process lends credence to the importance of neuronal connectivity in the mechanism for amyloid propagation in the human brain.

However, passive diffusion mechanisms only explain a portion of the variance observed, as the optimal NDM performance yielded moderate correlations. Notably, the diffusion mechanism models pathogenic agents as originating from a seed injection epicenter that then spreads throughout the system over time. The model does not take into account other external processes that may introduce protein aggregation, such as through metabolism and protein synthesis pathways [49]. We thus hypothesized that incorporation of such information into the NDM and increasing the pathological load systematically through a protein synthesis term may improve model performance.

### A. Extensions of the Network Diffusion Model

Recent work in the field has built upon the network diffusion model to derive the eNDM, where a first-order synthesis term at the seed node is included to model continuous synthesis of tau protein at the point of pathogenesis, in conjunction with the passive diffusion mechanism for spread [30]. In the present work, we have implemented the eNDM methodology into our modeling pipeline to examine amyloid pathophysiological spread, and found that the eNDM produced similar results to that of the baseline NDM. However, qualitatively the resulting modeled distribution does not recapture spatial locality as well as that of the baseline NDM. We attributed these shortcomings of the eNDM to an overly strict assumption where synthesis is hypothesized to only occur at the seed node, which does not reflect characteristics of wider-spread simultaneous accumulation of amyloid [52], [53].

As such, we expanded the eNDM to a definition with assumptions that more accurately reflected the biological hypotheses, expanding the synthesis term to instead apply to all nodes through our global eNDM, which we anticipated to improve upon baseline NDM performance. However, we found that applying the synthesis term in a global uniform manner resulted in performance that was indistinguishable from that of the baseline, yielding the same Pearson correlations. The result can be explained by the nature of the optimization. Since we are minimizing the SSE between spatial distributions of pathology and not the numerical pathological load at each region, the optimal distribution remains unaltered, though the synthesis-over-time introduces a delay as indicated by an increased optimal diffusion time. As such, applying global kinetics to an eNDM does not significantly impact the final model performance in the context of our optimization.

We thus surmise that both seed-based synthesis and globally uniform synthesis are two ends of a spectrum, and the reality may lie somewhere in between. A seed-based synthesis overly restricts protein pathogenesis via a synthesis process to a single location and is too specific, while global uniform synthesis with the same rate in all regions is too broad. Amyloid pathophysiology is hypothesized in the literature to be related to regional susceptibility, where some regions are found to be more vulnerable to pathological burden via properties such as cellular composition [54], molecular processes [55], and gene expression [56]. Global synthesis with a specific spatial pattern derived from regional vulnerability offers a potential middle ground between seed-based and uniform global synthesis, which we hypothesized would improve the model performance.

We sought to explore one dimension of regional vulnerability by modulating the rate parameter *λ* using relative regional genetic expression values *g*(*v*) from the AHBA in a gene eNDM framework, selecting genes that have been previously identified as significant genes in amyloid-*β* [49] as well as AD-related GWAS and PET-GWAS studies [50], [51]. When evaluating for performance improvements in the eNDM’s Pearson’s *R* metric over that of the baseline NDM, our results showed four gene eNDMs offer a significant improvement (Steiger’s Z *P*_FDR_ *<* 0.1) in the AD group when individually modulating *λ* with APOE, SORL1, FGL2, and IAPP regional expression values. The MCI group saw significant improvements from gene modulation with SORL1 and FGL2. Hence, we demonstrate that introducing regional vulnerability into the synthesis term, as defined through innate genetic expression from the AHBA, has offered an improvement over passive diffusion and may explain more of the variance observed in the ground truth amyloid PET pathology.

### B. Performance Gains in Gene eNDMs Reproduced in ADSP Cohort

After fitting the models in the ADNI, we froze model parameters and sought to predict the ADSP amyloid-*β* regional PET distributions. The models demonstrated good levels of generalizability, with moderate but significant correlation performance metrics that were slightly lower than those observed in the ADNI. Of note, three gene eNDM models (APOE, SORL1, and FGL2) exhibited significant performance improvements (Steiger’s Z, *P*_FDR_ *<* 0.10) over the baseline in the AD group. As such, we conclude that the inclusion of gene modulation and its improvements for these three genes are robust and generalizable.

The noted reduction in performance metrics compared to the ADNI group can be explained by a data limitation issue, where the PET imaging data obtained from the ADSP cohort did not have corresponding structural brain network data at the subject level. When freezing model parameters from the ADNI, we included the subpopulation brain networks from ADNI to resolve this limitation. In this regard, it can be said that the ADSP cohort experiences a low information scenario. Even so, our NDMs and eNDMs are able to still achieve significant Pearson’s correlation performance metrics (*P*_FDR_ *<* 0.01) and reproduce a number of gene contributions to model improvements, lending credence to the model’s robustness in a multi-cohort setting.

### C. Role of Genes That Improve eNDM Performance

The three genes that were observed to significantly improve model performance in both the ADNI and ADPS cohorts have also exhibited significant relationships to AD-related processes in previous studies.

APOE is noted as a risk gene in Alzheimer’s disease, particularly with the APOE-*ε*4 allele [57], [58], and its importance in improving the AD group eNDM performance was highly anticipated. We do note that the AHBA data does not specify the allele of APOE and what variant each subject in the dataset is carrying. In literature, *ε*3 and *ε*4 alleles are risk variants, while *ε*2 is potentially protective [59], which may be further explored in the eNDM framework with future disease-specific spatial genomics experiments.

SORL1 has been found to be an AD risk gene, participating in endocytotic amyloid-*β* trafficking mechanisms and dysfunction in microglial function [60], [61] and involved in recycling functions for neurons in the human connectome [62]. Deficiency in SORL1 has been shown to lead to decreased lysosomal activity, which in turn leads to impaired recycling and clearance rates of proteinopathies. Moreover, the impact of SORL1 on cellular processes is found to be tissue-specific, with increase vulnerability in the trans-entorhinal cortex regions [63]. In conjunction with our findings that region-specific modulation of gene eNDMs through SORL1 significantly improves model performance in MCI and AD groups, there is ample evidence that the gene plays an important role in the regional vulnerability characteristics that may lead to AD pathogenesis.

Lastly, FGL2 has been observed to be significantly associated to amyloid-*β* deposition in a previous PET-GWAS study, where participants that carried a minor allele associated with reduced FGL2 expression also exhibited increased amyloid-*β* deposition [64]. The gene’s role in amyloid is hypothesized to be neuroinflammation-related based on its role in immuno-suppressive functions in brain tumors [65], and may play a part in inflammation-related mechanisms such as amyloid-*β* clearance. The present study suggests a region-specific pattern as well, where innate regional vulnerability improves prediction of amyloid-*β* deposition patterns. Nevertheless, the gene is currently understudied in the context of AD, with increasing evidence of its role in the disease.

Our results illustrated that the inclusion of these genes as a regional vulnerability measure significantly improved the performance of a dynamic process in network diffusion, indicating a potential key role of these genes in amyloid-*β* spread. Further investigation of these genes may elucidate how genetic risk factors may lead to the downstream development of neurodegenerative disease.

### D. Introduction of Gene Modulation Increases Disease Specificity

In the gene eNDMs, it is observed systematically that introducing gene modulation offers a positive impact in disease groups, but no improvement or worsened results in healthy control. we saw that the Pearson correlation performance in the HC group generally decreased compared to baseline after introducing gene modulation, in both the ADNI and ADSP cohorts. In MCI, the performance differences were mixed, while in AD gene eNDMs exhibited performance increases. We interpret these results as regional genetic vulnerability driving a highly disease-specific spatial pattern, whereby spatial distributions of amyloid-*β* deposition in disease groups align more with spatial regional vulnerability than those of the healthy group. HC group is expected to have noisier, more random off-target binding of amyloid due to the process of normal aging [3], whereas AD subjects are expected to have more specific patterns. The gene eNDMs thus exhibit higher specificity towards disease group spatial patterns.

### E. Limitations

The present study has some limitations. ADSP was noted to lack corresponding structural brain network data, which was interpreted as the reason for lowered NDM and eNDM performance in the ADPS compared to ADNI. Spatial transcriptomics are obtained from AHBA which is a healthy subject cohort with only 6 subjects, with subcortical region data that is imputed. At present, this cohort is highly putative and accepted as a computational biology resource for spatial transcriptomics. In the future, when disease cohort spatial transcriptomics or larger datasets that maintain a similar level of high spatial specificity with more statistical power are available, they may improve the signal-to-noise ratio and identify impactful genes with greater confidence.

## V. Conclusion

In this work, we have applied NDM and eNDMs using white matter structural brain networks to model and characterize the spread of amyloid throughout the human brain. Our models were able to successfully predict and recapture ground truth patterns of amyloid deposition observed through FBP PET neuroimaging, for both the ADNI and ADSP cohorts. The involvement of the white matter brain network as the backbone for the diffusion process in the NDM lends credence to the hypothesis of trans-synaptic spread of amyloid in the human brain. Furthermore, inclusion of novel gene modulation in the eNDM framework to adjust the rate parameter using genetic regional vulnerability measures demonstrated marked improvements to eNDM performance, indicating the importance of regional vulnerability in predicting amyloid deposition patterns and impacts of genetic expression on pathophysiological processes in the brain. Future studies may further investigate disease-specific regional genomics in the context of eNDMs, and investigate individual-level predictions of pathophysiological phenomenon in the context of brain fingerprinting [40], [66]–[75].

## Acknowledgments

This work was supported in part by the National Institutes of Health grants T32 AG076411, F31 AG091902, RF1 AG068191, U01 AG066833, U01 AG068057, U19 AG074879, and R01 AG071470.

Data collection and sharing for this project were funded by the Alzheimer’s Disease Neuroimaging Initiative (ADNI) (National Institutes of Health Grant U01 AG024904) and DOD ADNI (Department of Defense award number W81XWH-12-2-0012). ADNI is funded by the National Institute on Aging, the National Institute of Biomedical Imaging and Bioengineering, and through generous contributions from the following: AbbVie, Alzheimer’s Association; Alzheimer’s Drug Discovery Foundation; Araclon Biotech; BioClinica, Inc.; Biogen; Bristol-Myers Squibb Company; CereSpir, Inc.; Cogstate; Eisai Inc.; Elan Pharmaceuticals, Inc.; Eli Lilly and Company; EuroImmun; F. Hoffmann-La Roche Ltd and its affiliated company Genentech, Inc.; Fujirebio; GE Healthcare; IXICO Ltd.; Janssen Alzheimer Immunotherapy Research & Development, LLC.; Johnson & Johnson Pharmaceutical Research & Development LLC.; Lumosity; Lundbeck; Merck & Co., Inc.; Meso Scale Diagnostics, LLC.; NeuroRx Research; Neurotrack Technologies; Novartis Pharmaceuticals Corporation; Pfizer Inc.; Piramal Imaging; Servier; Takeda Pharmaceutical Company; and Transition Therapeutics. The Canadian Institutes of Health Research is providing funds to support ADNI clinical sites in Canada. Private sector contributions are facilitated by the Foundation for the National Institutes of Health (www.fnih.org). The grantee organization is the Northern California Institute for Research and Education, and the study is coordinated by the Alzheimer’s Therapeutic Research Institute at the University of Southern California. ADNI data are disseminated by the Laboratory for Neuro Imaging at the University of Southern California.

ADSP data for this study were prepared, archived, and distributed by the National Institute on Aging Alzheimer’s Disease Data Storage Site (NIAGADS) at the University of Pennsylvania (U24-AG041689), funded by the National Institute on Aging.

## Notes

### Competing Interest Statement

The authors have declared no competing interest.

